# The *Pseudomonas protegens* PBL3 secretome exerts antimicrobial activity against *Burkholderia glumae* by disrupting cell envelope homeostasis and impairing bacterial replication

**DOI:** 10.64898/2026.11.04.764055

**Authors:** Biwesh Ojha, Dirk Anderson, Bara Altartouri, Gustavo Husein, Clemencia M. Rojas

**Affiliations:** Complex Biosystem Program, University of Nebraska-Lincoln; Department of Plant Pathology and Centre for Plant Science Innovation, University of Nebraska-Lincoln; Nebraska Centre for Biotechnology, University of Nebraska-Lincoln; Department of Biochemistry, University of Nebraska-Lincoln; Department of Genetics, College of Agriculture “Luiz de Queiroz” (ESALQ) University of São Paulo, Piracicaba, Brazil

## Abstract

The effectiveness of antimicrobials used to manage bacterial infections is increasingly threatened by the emergence of antimicrobial resistance. Many conventional antimicrobials act on a single defined target that pathogens can circumvent through target modification or pathway adaptation. In contrast, antimicrobials that disrupt multiple cellular processes may impose greater fitness costs and reduce the likelihood of resistance development, making their discovery a priority for pharmaceutical and agricultural applications. We previously demonstrated that the environmental bacterium *Pseudomonas protegens* PBL3 secretes a complex mixture of molecules (secretome) with antimicrobial activity against several plants and human pathogens. However, the mechanism underlying this activity remained unknown. Here, we used the plant pathogen *Burkholderia glumae* to investigate the mode of action of the *P. protegens* PBL3 secretome. We showed that the *P. protegens* PLB3 secretome causes extensive cell envelope damage that results in cell death and interferes with bacterial multiplication. Transcriptomic analysis further revealed broad disruption of processes associated with cell-envelope biogenesis and homeostasis and bacterial replication. Collectively, these findings demonstrate that the *P. protegens* PBL3 secretome acts as a multi-target antimicrobial that could be harnessed in the development of therapeutics to control bacterial infections of agricultural and medical importance.

## Introduction

Antimicrobials are widely used for the management of bacterial infections in humans, animals, and plants, yet their long-term effectiveness is increasingly challenged by the emergence of antimicrobial resistance (Hutchings et al., 2019). Antimicrobials inhibit growth or kill bacterial cells by interfering with essential bacterial processes such as cell wall synthesis, DNA replication, transcription or translation, by disrupting membrane integrity, or by altering essential metabolic pathways (Halawa et al., 2023; Kohanski et al., 2010).

Historically, antimicrobial discovery has often prioritized single-target antimicrobials with the rationale that a single target improves efficacy and reduces off-target effects; however, an important limitation of single-target antimicrobials is that mutations or modifications of the specific target, or of the pathways affected by a given antimicrobial lead to resistance, rendering it ineffective (Reygaert, 2018; Wenzel, 2020). In contrast, multi-target antimicrobials can affect several pathways simultaneously or target multiple cellular structures; by operating against multiple targets, these antimicrobials reduce the likelihood of resistance development as high fitness cost makes it unlikely that bacteria will acquire simultaneous resistance against multiple independent targets (Feng et al., 2023; Gray & Wenzel, 2020; Silver, 2007). The effectiveness of multi-target antimicrobials has driven the shift on drug discovery to curb the impact of bacterial infections (Feng et al., 2023; Wenzel, 2020).

Multi-target antimicrobials can be obtained from naturally occurring microorganisms that produce complex antimicrobial mixtures (multi-component) of structurally unrelated and functionally complementary compounds that simultaneously target different bacterial processes (Hutchings et al., 2019; Katz & Baltz, 2016; Talapko et al., 2022). Bacteria from the genera *Bacillus, Paenibacillus, Pseudomonas*, and *Streptomyces* are especially renowned for producing a diversity of metabolites that could be harnessed as sources of multi-target antimicrobials (Hutchings et al., 2019; Katz & Baltz, 2016). Accordingly, we previously showed that the environmental bacterium, *Pseudomonas protegens* PBL3, secretes a collection of molecules (secretome) with antimicrobial activity against pathogenic bacteria causing diseases in plants and humans (Ojha et al., 2025; Ortega et al., 2020). Although the specific antimicrobial molecules in the *P. protegens* PBL3 secretome remain to be identified, its chemical diversity suggests multi-component and multi-target activities that need to be dissected to further advance its use in the development as effective therapeutics.

In this study, we leveraged the plant pathogenic bacterium *Burkholderia glumae* as a model to investigate the cellular effects of the *P. protegens* PBL3 secretome. By combining live-cell imaging, flow cytometry and electron microscopy, we discovered that the *P. protegens* PBL3 secretome has pleiotropic effects including dramatic structural damage to the cell envelope leading to cell death, in addition to impairing bacterial replication. Through RNA sequencing, we further showed that these phenotypes are associated with the dysregulation of key pathways involved in cell envelope synthesis and maintenance, DNA replication, and cell division. Together, these findings provide mechanistic insight into the multi-target antimicrobial activity of the *P. protegens* PBL3 secretome to advance its further development as a source of effective therapeutics in agriculture and medicine.

## Materials and Methods

### Bacterial strains

*Burkholderia glumae* UAPB 13 and *P. protegens* PBL3 were streaked from-80 °C freezer on King’s B (KB) medium (King et al., 1954) and Luria Bertani (LB) agar plates, respectively and incubated at 28 °C for 2 days; a single colony was transfer to liquid medium for experiments.

*Burkholderia glumae* (*pGFP*) was generated by transforming *B. glumae* UAPB 13 with plasmid *pGFP-TIR* (Rojas et al., 2002) and transformants were selected and propagated on KB medium supplemented with gentamycin at a final concentration of 10 µg/mL.

### Preparation of *P. protegens* PBL3 secretome

*Pseudomonas protegens* PBL3 was grown in M9 minimal medium (1× M9 Minimal Salts [Sigma-Aldrich, St. Louis, MO, U.S.A.], 2 mM MgSO_4_·7H_2_O, 56 mM myo-inositol, and 0.2 mM CaCl_2_) in a fermentation tank to an optical density at 600 nm (OD_600_=0.5); the secreted fraction (secretome) was separated from whole cells by centrifugation, filter sterilization and lyophilization as previously described (Dahal et al., 2025).

### Time course growth assay

An overnight culture of *B. glumae* (*pGFP*) was adjusted to an OD_600_ of 0.2 (1 x 10^8^CFU/mL) with KB broth in a total volume of 4 mL and mixed with 1 mL of either M9 minimal medium or the *P. protegens* PBL3 secretome, each at 0.25 g/mL, and grown for 24 h at 28°C and constant agitation at 200 rpm. One hundred microliter aliquots were taken every 2 h for 24 h to measure OD_600_.

### Bacterial growth for cell viability assays

The effects of the *P. protegens* PBL3 secretome on *B. glumae* were evaluated at lag and log phases. To evaluate the effects of the *P. protegens* PBL3 secretome starting at lag phase, *B. glumae* (*pGFP*) liquid cultures grown overnight were adjusted to an initial OD_600_ of 0.2, mixed with 100 µL of the *P. protegens* PBL3 secretome at 0.25 g/mL and incubated for 18 h at 28 °C and constant agitation at 200 rpm. To measure the effects of the *P. protegens* PBL3 secretome starting at log phase, *B. glumae* (*pGFP*) liquid cultures at an initial OD_600_ of 0.2 were grown for 12 h before adding the *P. protegens* PBL3 secretome at 0.25 g/mL and incubated for additional 6 h at 28 °C and 200 rpm. Untreated *B. glumae* (*pGFP*) resuspended in water was used as a control for live cells, whereas *B. glumae* (*pGFP*) exposed to heat treatment (70 °C for 5 min) was used as a control for dead cells. Cell viability was evaluated by laser scanning confocal microscopy and flow cytometry.

### Confocal microscopy imaging

*Burkholderia glumae* (*pGFP*) cultures treated with the *P. protegens* PBL3 secretome either at the start of lag phase or at the start of log phase were centrifuged at 3,000 rpm for 15 min, resuspended in 400 µL of water, and stained with 0.5 µL of propidium iodide (PI) (1 µg/mL) for 15 min at room temperature. Ten microliters of stained cells were mounted on a microscope slide for imaging on a Nikon A1 upright laser scanning confocal microscope at 60x water immersion and 1.2 objective numerical aperture. Fluorescence was imaged using laser lines set up at 488 nm excitation/500-550 nm emission to collect GFP signal, and 561 nm excitation/570-620 nm emission to collect PI signal. The same settings were used across all samples. Confocal imaging settings were calibrated using unstained live *B. glumae* (*pGFP*) cells to confirm GFP fluorescence and heat-treated PI-stained cells to confirm red fluorescence from dead or membrane-compromised cells (Supplementary Figure S1A)

To quantify the images obtained from confocal microscopy, we imported images into Fiji (ImageJ) (Schindelin et al., 2012) and each confocal image was converted into an 8-bit image, with the threshold adjusted to observe bacterial cells only. After adjusting the threshold, the image was changed into binary mask, and the background was masked. The image was then watersheded to separate clustered bacterial cells into individual cells and enable accurate quantification of bacterial cells. Each image was then analyzed using the function ‘analysed particle’ to obtain the total number of bacterial cells within the image. Five independent images were analyzed in ImageJ to quantify bacterial counts, and the mean values with standard deviations were calculated.

### Flow cytometry analysis

*Burkholderia glumae* (*pGFP*) treated with the *P. protegens* PBL3 secretome and untreated control were stained with PI at room temperature for 15 min before scanning. Flow cytometry settings were calibrated and gated using unstained *B. glumae* (*pGFP*) cells to define the live GFP-positive population and heat-treated *B. glumae* (*pGFP*) cells stained with propidium iodide (PI) to define the dead PI-positive population fluorescing in the red channel (Supplementary Figure S1B). After calibration, stained samples were loaded on CytoFLEX LX with CytExpert software (Beckman Coulter) and excited with 488 nm and 561 nm lasers to generate GFP and PI images, respectively. Image analysis was performed using FlowJo (BD Biosciences).

### Transmission Electron Microscopy (TEM) and Scanning Electron Microscopy (SEM)

*Burkholderia glumae* cultures treated with *P. protegens* PBL3 secretome were fixed using 2.5% glutaraldehyde for 1 h at room temperature for both SEM and TEM. After fixation, cell cultures were centrifuged at 3,500 rpm for 10 min to obtain the bacterial pellet, which was then resuspended and washed three times with 0.1 M sodium cacodylate (CA) buffer (sodium cacodylate salt and water, PH 7.2), followed by post-fixation with a 1% osmium tetroxide for 1 h at room temperature. Samples were then washed three times with CA buffer and dehydrated in a graded ethanol series (30%, 50%, 70%, 95%, 95%, 100%, and 100%) for 20 min each. For TEM after dehydration with 100% ethanol, samples were infiltrated with a 1:1 mixture of Spurr resin and ethanol twice for 3 h each, followed by three changes of 100% Spurr resin, with the last one left overnight at 4 °C. Polymerization was done at 65 °C for 24 h. Thin sections (80nm) of the samples were cut using a Leica EM UC7 ultramicrotome, which was then placed on copper grids and post-stained with 2% uranyl acetate, followed by Reynolds’ lead citrate. Sections were observed under Hitachi HT-7800 TEM at 80 KV.

For SEM, samples were mounted on a filter membrane on aluminum SEM stubs and left to dry at room temperature before being sputter-coated with chromium. Images were collected using a Hitachi FlexSEM 1000-II at 5 kV.

### RNA sequencing

*Burkholderia glumae* cells exposed to M9 minimal medium or the *P. protegens* PBL3 secretome and harvested after 4 and 8 h. Three biological replicates were included for each condition and time point. Total RNA was isolated using the RNeasy Mini Kit (QIAGEN, Germany) and residual genomic DNA was removed following standard procedures. The resulting RNA samples were submitted to University of California-Davis Genome Center DNA Technologies and Expression Analysis Core for library preparation and was sequenced on an Illumina NovaSeq X 25B (Illumina, San Diego, CA) with paired-end 150 bp reads.

Raw RNA-seq reads were initially assessed for quality using FastQC (v0.12.1) (Andrews, 2010). Adapter sequences and low-quality bases were removed using Trimmomatic (v0.40-rc1) (Bolger et al., 2014) with the following parameters: ILLUMINACLIP:TruSeq3-PE-2.fa:2:30:10, LEADING:3, TRAILING:3, SLIDINGWINDOW:4:15, and MINLEN:36. Post-trimming quality was re-evaluated to ensure the removal of technical artifacts. High-quality filtered reads were aligned to the *B. glumae* UAPB13 reference genome (NCBI RefSeq: GCF_024611525.1) using HISAT2 (v2.1.0) (Kim et al., 2015). Alignment files were converted to BAM format, sorted, and indexed using Samtools (v1.10) (Li et al., 2009). Gene-level quantification was performed using featureCounts from the Subread package (v2.0.0) (Liao et al., 2014), assigning reads to coding sequences based on the locus_tag attribute of the RefSeq GFF annotation. The resulting count matrix was then analyzed using DESeq2 (v1.40.2) (Love et al., 2014) to identify differentially expressed genes. To improve effect size estimation, log2 fold change values were shrunk using the adaptive shrinkage (ashr) method. Heatmaps were generated using the *pheatmap* package in R to visualize the expression pattern of genes across treatment conditions and time points.

## Results

### The *P. protegens* PBL3 secretome reduces bacterial populations by causing cell death

To define critical stages of *B. glumae’s* growth exhibiting sensitivity to the *P. protegens* PBL3 secretome, we monitored the growth of *B. glumae* (*pGFP)* in KB medium (volume adjusted with water), KB amended with M9 medium as control, or KB amended with the *P. protegens* PBL3 secretome over 24 h. At early time points (0–6 h), *B. glumae* (*pGFP*) cultures grown on KB, KB amended with M9 medium or KB amended with the *P. protegens* PBL3 secretome were in lag phase (Figure 1A). From 6 to 24 h, *B. glumae* (*pGFP*) cultures grown on KB or KB amended with M9 medium (Purple and Blue lines in graph) showed the typical exponential growth phase until reaching an OD_600_ value of 2.7 at 24 h. However, *B. glumae* (*pGFP*) grown on KB amended with the *P. protegens* PBL3 secretome (orange line in graph) was arrested in lag phase and remained at OD_600_ of 0.5 until 24 h, suggesting that the *P. protegens* PBL3 secretome either interferes with *B. glumae* (*pGFP*)’s growth or causes cell death.

**Figure 1.**
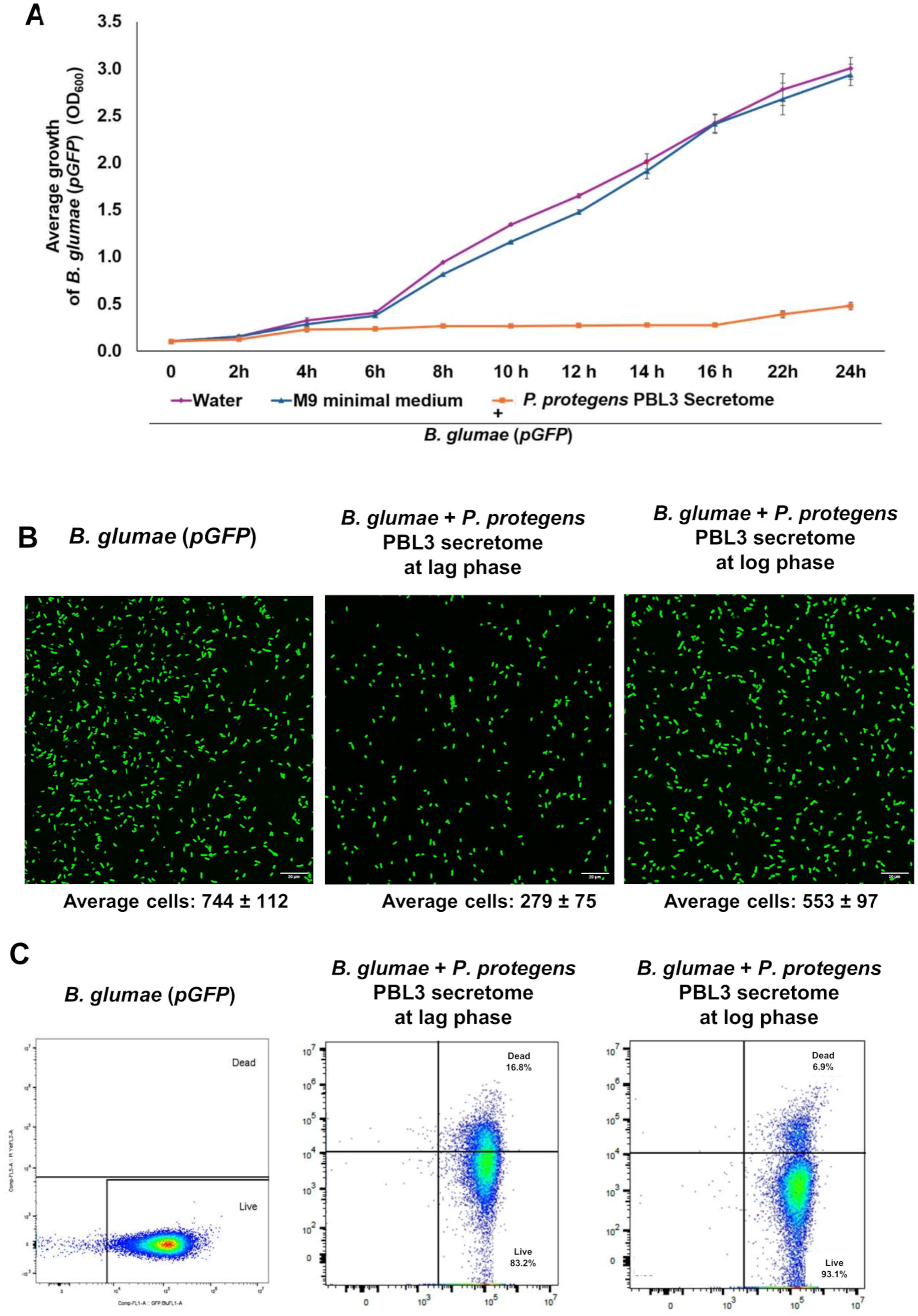
The *P. protegens* PBL3 secretome and increases membrane permeability causing bacterial cell death. **(A)** *B. glumae (pGFP)* was grown on KB broth mixed with either water (purple line), M9 minimal medium (blue line) or the *P. protegens* PBL3 secretome (orange line), and its growth evaluated as optical density at 600 nm (OD_600_) measured every 2 h for 24 h. **(B,C)** *B. glumae (pGFP)* mixed with M9 minimal medium or the *P. protegens* PBL3 secretome, added either at the start of the lag phase or at the start of the mid-log phase, was stained with propidium iodide to assess cell viability by confocal imaging **(B)** or flow cytometry **(C)**. Confocal images were obtained in green and red channels. Scale bar: 20µm.

To distinguish between cell death and growth inhibition, we incubated *B. glumae* (*pGFP*) with the *P. protegens* PBL3 secretome at the start of lag phase (to monitor the effect of the *P. protegens* PBL3 secretome on cell division), and at the start of log phase (to monitor its effect on cells that are actively growing), and then evaluated bacterial populations by laser scanning confocal microscopy. When comparing *B. glumae* (*pGFP*) fluorescence control and *B. glumae* (*pGFP*) fluorescence after treatment with the *P. protegens* PBL3 secretome, there was a 62.5% reduction in fluorescent bacteria when the *P. protegens* PBL3 was added at lag phase and a 26% reduction in fluorescent bacteria when *P. protegens* PBL3 secretome was added at mid-log phase (Figure 1B), indicating that bacterial cells at lag phase are more vulnerable to the *P. protegens* PBL3 secretome. This reduction in *B. glumae* (*pGFP*) fluorescence led us to hypothesize that the *P. protegens* PBL3 secretome causes cell death, and therefore we quantified cell viability by evaluating the internalization of the DNA-binding dye propidium iodide (PI) through flow cytometry (Davey & Guyot, 2020). Flow cytometry revealed that *B. glumae* (*pGFP*) control shows a pattern where most cells are fluorescent green indicating live cells. In contrast, treatment with the *P. protegens* PBL3 secretome caused a distinct shift of bacterial cell populations toward the PI-positive region, indicating that the *P. protegens* PBL3 secretome caused membrane permeabilization enabling the internalization of PI that accounted for 16.8% cell death when added at lag phase and 6.9% when added at the start of mid-log phase (Figure 1C).

### The *P. protegens* PBL3 secretome causes structural damage to the bacterial cell envelope

The permeabilization of bacterial cells associated with the *P. protegens* PBL3 treatment led us to hypothesize that one possible mechanism of action is damage to the *B. glumae’*s cell envelope. To gain deeper insight into the structural changes in the *B. glumae* due to *P. protegens* PBL3 secretome, we used transmission electron microscopy (TEM) to visualize *B. glumae* cells treated with M9 minimal medium (control) or with the *P. protegens* PBL3 secretome added at lag phase and incubated for 8 h, the time point at which there was a clear distinction between M9 minimal medium treatment and treatment with the *P. protegens* PBL3 secretome, as shown in Figure 1A. *B. glumae* treated with M9 minimal medium had a continuous cell envelope (Figure 2, green arrows), whereas *B. glumae* treated with *P. protegens* PBL3 secretome had visible breaks or gaps in the cell envelope resulting in the leakage of cytoplasmic content (Figure 2, blue arrow). More severe effects include cells completely devoid of cellular content (Figure 2, purple arrow) or with loss of defined membranes (Figure 2, black arrow). These dramatic disruptions in the cell envelope integrity account for ensuing cell death.

**Figure 2.**
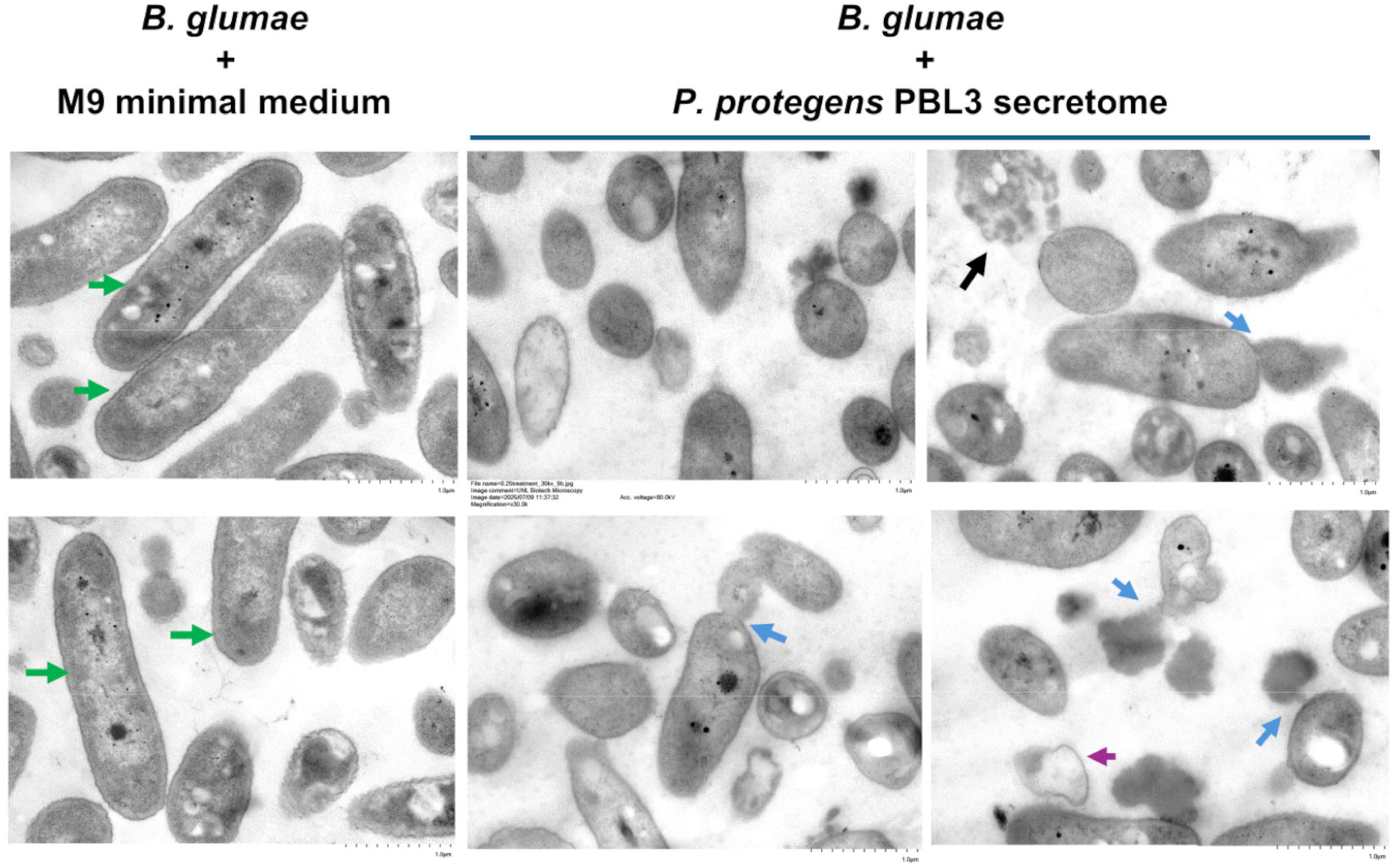
**The *P. protegens* PBL3 secretome disrupts *B. glumae*’s cell envelope.** *B. glumae* cultures grown in medium amended with M9 minimal medium (control) or the *P. protegens* PBL3 secretome at the start of lag phase for 8 h and processed for TEM and imaged. Color arrows show morphological differences between control cells and *P. protegens* PBL3-treated cells. Green arrows: intact envelope structure, Blue arrows: envelope rupture and leakage of cytoplasm, Black arrow: loss of defined membrane, Purple arrow: bacterial cells devoid of cytoplasm. (Scale bar=1 µm)

### The *P. protegens* PBL3 secretome inhibits bacterial replication and alters cell morphology

We demonstrated that adding the *P. protegens* PBL3 secretome at the start of lag phase caused a significant impact in bacterial populations, as evaluated by the bacterial growth curve, yet the percentage of cell death as quantified by flow cytometry was only 16.8%, suggesting that cell death alone did not account for the reduction in bacterial numbers. Thus, an alternative hypothesis is that the *P. protegens* PBL3 interferes with bacterial replication, which was tested by leveraging flow cytometry data on forward scatter light (FSC), a parameter that directly measures cell size (Bajgelman, 2019). At 8 h, *B. glumae* (*pGFP*) grown in culture amended with M9 medium (control) revealed that *B. glumae* (*pGFP*) cells exhibited a marked shift toward higher FSC-A values consistent with increased cell size and elongation, which are associated with replication (Figure 3A, left, top panel). In contrast, *B. glumae* (*pGFP*) grown in culture medium amended with the *P. protegens* PBL3 secretome showed cells clustered at lower FSC-A values, indicating the inhibition of cell elongation (Figure 3A, right top panel). Quantification of FSC-A values showed that *P. protegens* PBL3 secretome-treated cells had 51% lower FSC-A compared with the control at 8 h (Supplementary Figure 2A). At 18 h, cells in both groups reached stationary phase which stops bacterial replication and consequently, a reduction in FSC-A values was observed regardless of treatment conditions. Nevertheless, quantification of FSC-A values between control and *P. protegens* PBL3 treatment at this time point, showed about 5% lower FSC-A in *P. protegens* PBL3-treated cells than control treatment (Supplementary Figure 2A).

**Figure 3.**
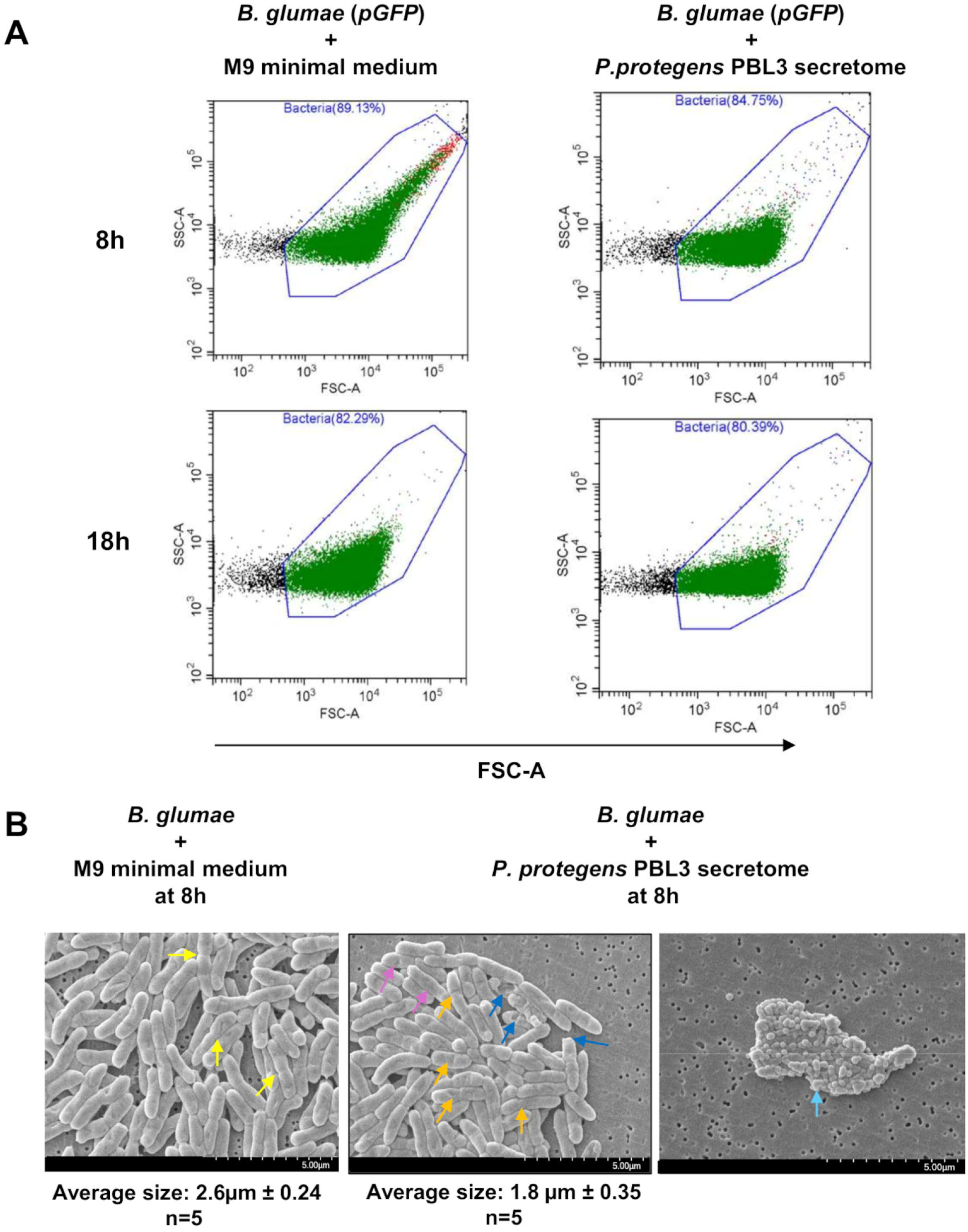
The *Pseudomonas protegens* PBL3 secretome Inhibits Cell Elongation and Replication of *B. glumae*. **(A)** Flow cytometry on *B. glumae (pGFP)* mixed with M9 minimal medium or the *P. protegens* PBL3 secretome added at the start of the lag phase and stained with propidium iodide was used to assess cell elongation and replication based on forward-scatter (FSC) and side-scatter (SSC) light signals. **(B)** *B. glumae (pGFP)* cultures were treated with the *P. protegens* PBL3 secretome at lag phase for 8h. Colored arrow showed differences in *B. glumae* cell morphology between control and *P. protegens* PBL3 secretome treatment. Yellow arrow: *B. glumae* cells undergoing replication, Purple: crevices and folding of *B. glumae* cells, Orange: Constriction and cell damage, Turqoise: change from rod to coccoid shape. (Scale bar= 5 µm)

To confirm the phenotypes of the *P. protegens* PBL3 secretome on cell elongation and size, we used scanning electron microscopy (SEM) of *B. glumae* control treatment versus treatment with the *P. protegens* PBL3 secretome. We initially examined cells after 18 h of *P. protegens* PBL3 treatment when added at the start of lag phase. However, at this point, it was not possible to obtain clear visualization of *B. glumae* cells given the abundant cellular debris (Supplementary Figure 2B, red arrows). We then chose only the 8 h time point for all experiments to better capture the morphological changes induced by the *P. protegens* PBL3 secretome before extensive cell disruption and debris accumulation occurred.

At 8 h, images of *B. glumae* with control treatment show the expected elongated rod shapes with average length of 2.6 µm, whereas the average size of *B. glumae* treated with the *P. protegens* PBL3 secretome was 1.8 µm. Control treatment also revealed bacterial cells undergoing cell division (Figure 3B, left panel, yellow arrow), that were not observed in *B. glumae* cells treated with the *P. protegens* PBL3 secretome (Figure 3B, middle panel). We also observed that *B. glumae* cells treated with *P. protegens* PBL3 secretome exhibited clear crevices in the cell envelope (Figure 3B, middle panel, blue color), folding (Figure 3B, middle panel, purple arrow), constriction (Figure 3B, middle panel, orange arrow), and in some cases, a transition to coccoid morphology (Figures 3B, right panel, turquoise arrow). This is important because the latter two phenotypes are associated with cell envelope integrity, which suggests that the *P. protegens* PBL3 secretome affects bacterial cell envelope composition.

### The *P. protegens* PBL3 secretome alter expression of genes associated with cell envelope synthesis, maintenance and cell shape maintenance

The envelope damage and altered bacterial morphology observed at 8 h after *P. protegens* PBL3 secretome treatment led us to evaluate transcriptional changes associated with the phenotypes observed. Since those phenotypic changes were observed at 8 h post-treatment, we posit transcriptional changes occur earlier. Thus, to identify early and progressive transcriptional responses, we performed RNA-seq analysis of *B. glumae* cells treated with the *P. protegens* PBL3 secretome or control medium at 4 h and 8 h after treatment. Principal component analysis (PCA) of RNA-seq expression profiles showed clear separation among *B. glumae* cells by both treatment and time point (Supplementary Figure S3A). PC1 explained 68% of the total variance between the controls and the treated samples, indicating a strong transcriptional effect of *P. protegens* PBL3 secretome exposure. PC2 explained an additional 25% of the variance between the 4 h and 8 h samples, suggesting time-dependent transcriptional changes. Biological replicates clustered tightly within each group, supporting the reproducibility of the RNA-seq dataset. Volcano plots revealed extensive differential gene expressions in *B. glumae* treated with the *P. protegens* PBL3 secretome at both 4 h and 8 h. Although substantial transcriptional changes were already evident at 4 h (858 DEGs), the broader distribution and greater absolute log_2_-fold changes at 8 h (1785 DEGs) indicated a progressive intensification of bacterial responses to secretome exposure (Supplementary Figure S3B).

The dramatic structural changes in the cell envelope led us to analyze transcriptional profiles of genes involved in processes occurring in the several layers of the cell envelope: outer membrane, the peptidoglycan cell wall and the inner membrane. Two major processes occurring at the outer membrane include the synthesis of lypopolysacharide (LPS) and the incorporation of outer membrane proteins (OMP) mediated by *lpx* and β-barrel assembly machine (*Bam*) genes, respectively. LPS assembly requires synthesis of lipid A mediated by *lpx* genes and additional genes for the core assembly. When comparing expression of those genes between *B. glumae* treated with the *P. protegens* PBL3 secretome and control, we found that *lpxA*, *lpxB*, *lpxK*, *lpxD*, and *lpxL* showed no significant change in expression at 4 h but were consistently downregulated at 8 h, whereas some genes that participate in the core assembly such as *rfaE1*, *kdsA*, and *gmhB*, were significantly induced at 4 h but were repressed at 8 h. Intriguingly D-glycero-alpha-D-manno-heptose-1,7-bisphosphate 7-phosphatase (*JWI72_RS01505*) that provides sugars for the building of the LPS showed a unique pattern of expression, with no significant differences in expression between *P. protegens* PBL3 treatment and control at 4 h but with strong induction with the *P. protegens* PBL3 treatment at 8 h (Figure 4A).

**Figure 4.**
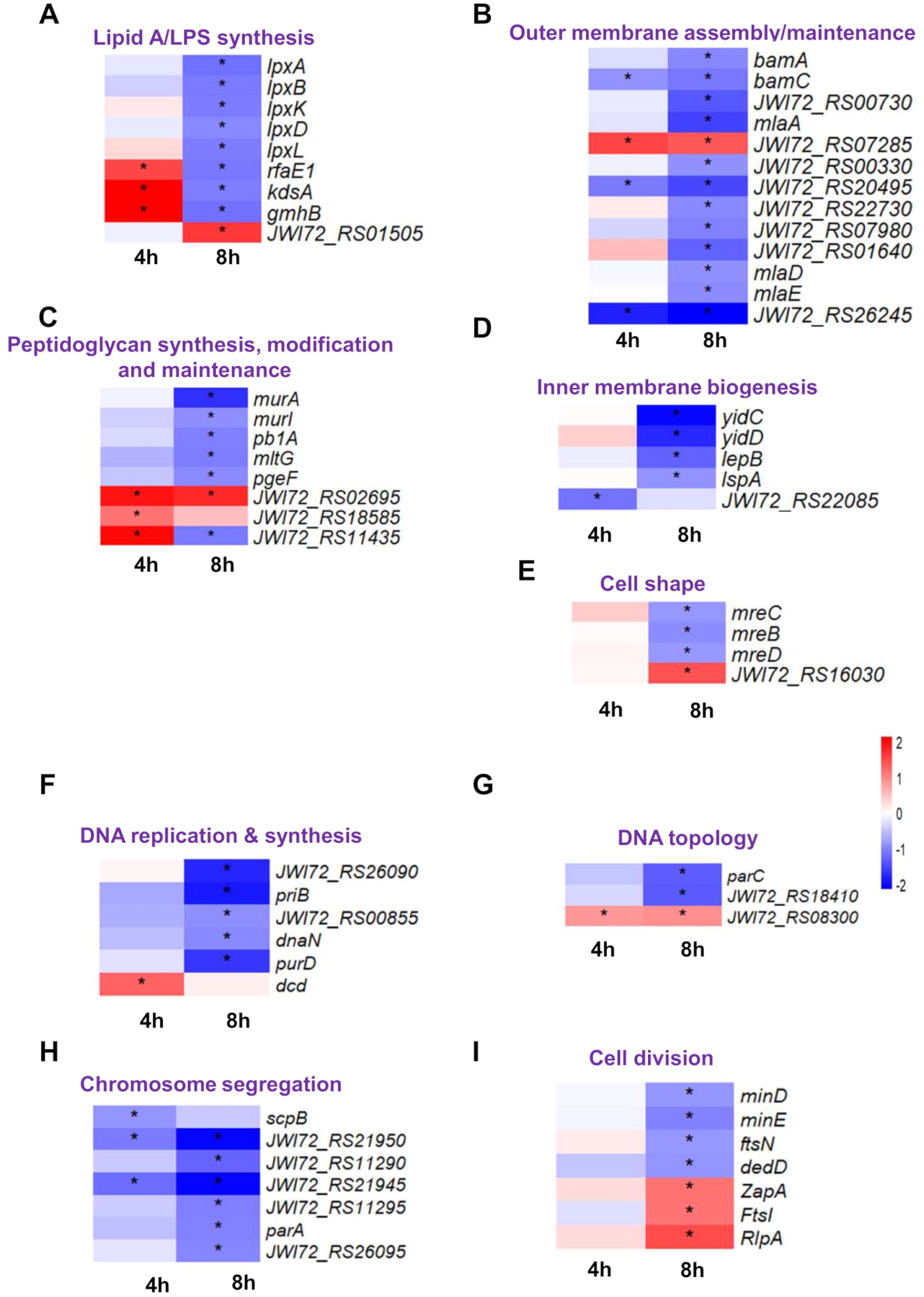
The *Pseudomonas protegens* PBL3 alter the expression of genes required for cell envelope composition and bacterial replication. Transcriptomic changes in *B. glumae* after treatment with the *P. protegens* PBL3 secretome were evaluated after 4 h and 8 h post-treatment and in comparison, with M9 medium controls. Heatmaps show Differentially Expressed Genes in selected functional categories including cell envelope synthesis and maintenance (A-E), and bacterial replication (F-I) Red indicates upregulation and blue color indicates downregulation relative to the corresponding control. Asterisks within heatmap cells indicate genes that were significantly differentially expressed relative to the control.

Among *Bam* genes, treatment with the *P. protegens* PBL3 secretome caused early repression of *bamC* at 4 h and both *bamA* and *bamC* were downregulated at 8h. Other genes involved in outer membrane biogenesis and maintenance also showed disregulation associated with *P. protegens* PBL3 secretome treatment. SurA N-terminal domain-containing protein (*JWI72_RS00730*), encoding a periplasmic chaperone, genes encoding several OMPs, such as OmpH (*JWI72_RS00330*), OmpW/AlkL family proteins (*JWI72_RS22730* and *JWI72_RS01640*), and MipA/OmpV family protein (*JWI72_RS07980*), as well as genes associated with the Mla lipid asymmetry maintenance pathway, including *mlaA*, *mlaD*, and *mlaE* were downregulated only 8 h after *P. protegens* PBL3 secretome treatment in comparison with control, whereas the genes encoding OmpA family protein (*JWI72_RS20495*) and the outer membrane lipoprotein-sorting protein (*JWI72_RS26245*) were downregulated at both 4 and 8 h; and only the gene encoding an OmpA/MotB family protein (*JWI72_RS07285*), was induced at both time points (Figure 4B).

We also found a disrupted and time-dependent expression pattern among genes involved in peptidoglycan synthesis, modification, and remodeling (Figure 4C). At 4 h, genes involved in peptidoglycan precursor synthesis, *murA* and *murl*, as well as genes associated with peptidoglycan polymerization and remodeling, including penicillin-binding protein 1A (*pbp1A*), endolytic transglycosylase (*mltG*), and the peptidoglycan editing factor (*pgeF*), showed no significant differential expression between the control and secretome-treated cells. However, at 8 h, all those genes were significantly downregulated following treatment with the *P. protegens* PBL3 secretome. In contrast, the genes involved in peptidoglycan remodeling, M23 family metallopeptidase (*JWI72_RS02695*) and an aspartate/glutamate racemase family protein (*JWI72_RS18585*) were strongly induced at 4 h; however, only the M23 family metallopeptidase remained induced at 8 h. A lytic transglycosylase domain-containing protein (*JWI72_RS11435*) was induced at 4 h but repressed by 8 h (Figure 4C).

The *P. protegens* PBL3 secretome treatment also affected genes involved in inner membrane protein biogenesis (Figure 4D). Genes encoding the membrane protein insertase (y*idC*), membrane insertion efficiency factor (*yidD*), signal peptidase I (*lepB*), and lipoprotein signal peptidase (*lspA*) showed no significant differential expression at 4 h but were strongly repressed at 8 h. In contrast, the gene encoding YihY family inner membrane protein (*JWI72_RS22085*), which is a conserved bacterial protein involved in transport of inner proteins was repressed at 4 h but it was not differentially expressed at 8 h.

Because treatment with the *P. protegens* PBL3 secretome also caused altered bacterial morphology, we also examined *mre* genes associated with rod-shape maintenance (Figure 4E) and found that genes encoding core components of the Mre system, *mreB*, *mreC*, and *mreD,* showed no significant differential expression at 4 h but were significantly repressed by 8 h. In contrast, another rod shape-determining protein (*JWI72_RS16030*), was induced at 8 h (Figure 4E).

These results show that bacterial exposure to the *P. protegens* PBL3 secretome results in broadly transcriptomics changes in different pathways required for cell envelope formation, with the expression of few genes affected as early as 4 h post treatment, but the majority of changes observed at 8 h post-treatment, which are consistent with cell-envelope damage and altered cellular morphology observed.

### The *P. protegens* PBL3 secretome alters the expression of genes associated with cell division and DNA replication

Our observations that the *P. protegens* PBL3 secretome not only causes cell death but interfere with bacterial cell division, prompted us to evaluate the expression of genes related to bacterial replication and cell division. When evaluating genes associated with DNA replication and synthesis, we observed that several genes were significantly repressed at 8 h after treatment with the *P. protegens* PBL3 secretome in comparison with M9 medium control treatment and they included *JWI72_RS26090,* encoding the replication initiation protein, *priB* encoding the primosomal replication protein, as well as genes encoding three DNA polymerase III-associated components *JWI72_RS00855, dnaN* encoding DNA polymerase III subunit beta and *purD* encoding phosphoribosylamine--glycine ligase. In contrast, *dcd* encoding dCTP deaminase was strongly induced at 4 h but was not significantly differentially expressed by 8 h (Figure 4F).

Genes associated with DNA topology also showed a time-dependent transcriptional response following exposure to the *P. protegens* PBL3 secretome (Figure 4G). Genes encoding DNA topoisomerase IV-associated subunits, including *parC* and *JWI72_RS18410*, did not show differential expression between control and *P. protegens* PBL3 secretome treatment at 4 h but were strongly repressed at 8 h. In contrast, the gene encoding DNA topoisomerase IB (*JWI72_RS08300*) was induced at both 4 h and 8 h.

Some genes associated with chromosome segregation were also repressed following treatment with the *P. protegens* PBL3 secretome at 4 h such as *scpB*, encoding the SMC-Scp complex subunit, as well as *JWI72_RS21950* and *JWI72_RS21945* encoding ParB/RepB/Spo0J-family partition protein and ParA family protein, respectively. The latter two genes showed higher levels of repression at 8h, whereas *scpB* levels of expression at 8h were not significantly different between *P. protegens* PBL3 secreteome treatment and control. Additional genes also belonging to ParA family proteins (*JWI72_RS11295* and *parA*) and ParB/RepB/Spo0J family partition proteins (*JWI72_RS11290* and *JWI72_RS26095*) were not significantly differentially expressed at 4 h but were repressed at 8 h (Figure 4H).

Genes associated with cell division showed no differential expression when comparing *P. protegens* PBL3 and control treatments at 4 h. However, by 8 h, *P. protegens* PBL3 secretome treatment caused downregulation of genes involved in septum site selection such as *minD* and *minE*, as well as genes encoding SPOR domain-containing division proteins, including *ftsN* and *dedD*. In contrast, several genes were induced at 8h, including zapA, a conserved gene related to cell division, as well as genes encoding a transpeptidase (*FtsI*) and family septal ring lytic transglycosylase (*RlpA*).

These results show that exposure to the *P. protegens* PBL3 secretome progressively alters the expression of genes associated with DNA replication, chromosome organization, and cell division, with the strongest transcriptional changes occurring at 8 h. These changes are consistent with the growth arrest, impaired cell division, and altered cellular morphology observed in *B. glumae* following secretome treatment.

## Discussion

The *P. protegens* PBL3 secretome is expected to be a complex mixture of molecules based on the identification of at least 14 putative biosynthetic gene clusters (BGCs) predicted to encode secondary metabolites, some of them already recognized for their antimicrobial activity (Ortega et al., 2020). Moreover, because some of these metabolites are predicted to be generated through non-ribosomal synthesis, a single BGC can generate a diversity of molecules that cannot be predicted from BGC sequences alone (Butler et al., 2026; Iacovelli et al., 2021; Patil & Gore, 2025). Thus, the quantity and diversity of potential antimicrobial molecules in the *P. protegens* PBL3 secretome are anticipated to have different targets and mechanisms of action, collectively contributing to a robust outcome. Accordingly, in this study, we showed that the *P. protegens* PBL3 secretome have pleiotropic effects disrupting the structural integrity of the cell envelope and altering bacterial replication, each of these processes having a detrimental effect on bacterial viability.

As a Gram-negative bacterium, the *B. glumae’*s envelope comprises the inner membrane, the peptidoglycan layer, and the outer membrane, which function together to maintain cellular integrity and morphology (Delcour, 2009; Silhavy et al., 2010). This composition makes the Gram-negative cell envelope a powerful barrier that prevents antibiotic penetration hindering many antimicrobial therapies. Thus, targeting this structure represents an effective strategy to control bacterial infections (Ntallis et al., 2026). Our combined confocal microscopy, flow cytometry, SEM and TEM data provided compelling evidence that the *P. protegens* PBL3 secretome causes dramatic structural damage to the cell envelope leading to leakage of cytoplasmic contents and subsequent cell death. Previously, the use of a crude extract from *P. protegens* Pf-5 also caused crevices and morphological changes in the plant pathogen *Pantoea ananatis* as shown by SEM and TEM imaging (Gu et al., 2022), suggesting that common metabolites between *P. protegens* Pf-5 and *P. protegens* PBL3 could be responsible for the disruption of the bacterial cell envelope. These two strains have as common metabolites 2,4-diacetylphloroglucinol and pyoluteorin (Dahal et al., 2024; Gu et al., 2022), which were previously shown to cause membrane permeabilization and profound structural changes in *Escherichia coli* and *Pseudomonas aeruginosa* PAO1, respectively (Gu et al., 2022; Julian et al., 2021). However, when we previously tested purified commercially available 2,4 DAPG and pyoluteorin, we showed that they did not significantly contribute to the antimicrobial activity against *B. glumae* (Dahal et al., 2024), making it unlikely that these molecules are responsible for the phenotypes observed.

While multiple components of the Gram-negative bacteria cell envelope are targets of antimicrobials (Ntallis et al., 2026), for example polymyxins targeting LPS and Darobactin targeting insertion of OMP (Ntallis et al., 2026; Page & Walker, 2021), our data does not define the specific molecules in the *P. protegens* PBL3 secretome causing changes in the cell envelope, nor defines which components of the *B. glumae* envelope are targeted by the *P. protegens* PBL3 secretome. Rather, our transcriptomics data highlights that the dramatic structural changes in the cell envelope are associated with alteration of multiple pathways responsible for biosynthesis and maintenance of the different layers of the cell envelope.

The outer membrane includes the LPS layer whose biosynthesis requires the assembly of lipid A and the core oligosaccharides and we showed that the *P. protegens* PBL3 secretome downregulates *lpx A*, *lpxB*, *lpxK*, *lpxD* and *lpxL*, required for lipid A biosynthesis (Raetz et al., 2009), as well as genes such as *rfaE1, gmhB* and *kdsA* that function in assembly of the oligosaccharide core. Besides the LPS layer, the outer membrane also has a glycerophospholipid layer with embedded OMP, which are inserted through the activities of BamA and SurA (Fenn et al., 2024), and our transcriptomics data showed that the *P. protegens* PBL3 secretome not only downregulates *BamA* and *SurA* but also several genes encoding several OMP proteins such as OmpA, OmpH, OmpV and OmpW. Additional proteins localized to the outer membrane include proteins from the Mla system that are responsible for maintaining the balance of glycerophospholipids at the outer membrane {Kaur, 2024 #209} and genes encoding several of those proteins were also downregulated by the *P. protegens* PBL3 secretome. These transcriptional changes suggest that the secretome may compromise multiple processes required for outer membrane assembly and maintenance. Besides the effects on the outer membrane, the *P. protegens* PBL3 secretome also caused downregulation of genes involved in peptidoglycan synthesis and maintenance such as *murA* and *murI*, *pb1A, mltG*, and *pgeF* as well as genes participating in the biogenesis of the inner membrane such as *yidC, yidD, lepB,* and *lspA* suggesting a possible imbalance between cell envelope synthesis and remodeling. While this transcriptomic data uncovered specific genes affected by the *P. protegens* PBL3 secretome, it is not clear if the secretome directly alters their expression or whether the structural damage caused by the secretome triggers a collective stress response to shut down essential processes, while the induction of few genes might occur to repair damage. A general shut down is consistent with the downregulation of the *Mre* genes controlling cell shape which resulted in the transition from rod to cocci shape.

In addition to affecting cell envelope biogenesis and maintenance, the *P. protegens* PBL3 secretome also altered the expression of genes involved in bacterial multiplication, which requires coordinated DNA replication, chromosome segregation, and cell division (O’Donnell et al., 2013). DNA replication involves proteins responsible for replication initiation, restart, and DNA synthesis, and our transcriptomic data showed that the secretome downregulated genes encoding proteins associated with these functions, as well as *purD*, involved in purine biosynthesis. Following DNA replication, bacterial chromosomes must be properly organized and segregated before cell division, a process involving topoisomerase IV, which resolves interlinked daughter chromosomes, and ParA/ParB-family proteins, which contribute to chromosome organization and segregation (O’Donnell et al., 2013; Reyes-Lamothe & Sherratt, 2019). Genes encoding these proteins were downregulated following secretome treatment, whereas a gene encoding topoisomerase IB, involved in regulating DNA topology, was induced, although the significance of this induction remains unclear. Cell division requires proper positioning and assembly of the divisome, followed by septal peptidoglycan synthesis and envelope constriction to separate daughter cells. The Min system, including *MinD* and *MinE,* regulates division-site positioning, while *ZapA* contributes to FtsZ-ring organization*. FtsN* participates in septal constriction, whereas *FtsI* and *RlpA* contribute to septal peptidoglycan synthesis and remodeling, respectively (Mahone & Goley, 2020). We found that the *P. protegens* PBL3 secretome downregulated *minD*, *minE*, and *ftsN*, while *zapA* and genes encoding FtsI-family and RlpA proteins were induced. Although these findings indicate altered expression of genes involved in multiple processes required for bacterial multiplication, it remains unclear whether these changes reflect direct effects of the secretome or secondary responses to cellular damage. The broad downregulation of genes involved in DNA replication, chromosome segregation, and cell division is consistent with the observed growth inhibition, whereas induction of selected genes may represent compensatory responses to disrupted cellular processes.

Overall, our study demonstrates that the *P. protegens* PBL3 secretome exerts antimicrobial activity against *B. glumae* by disrupting cell-envelope integrity and affecting multiple essential cellular processes associated with bacterial growth and proliferation. The multiple transcriptomic changes and phenotypes observed associated with *P. protegens* PBL3 treatment are consistent with a multi-component/ multi-target antimicrobial. These findings support the PBL3 secretome as a source of bioactive molecules with multi-process antimicrobial activity and potential for the development of new antimicrobial strategies.

## Acknowledgments

We would like to thank Connor Pederson, Nebraska Centre for Biotechnology, for his assistance with TEM and SEM imaging. We would like to thank Thi Kim Ha Nguyen and Shilu Dahal for critically reviewing the manuscript. This work was supported by Nebraska Research Initiatives and National Science Foundation (Award no. 2334721)

## Author Contributions

BO: Performed experiments, analyzed data, prepared and edited manuscript; DA, BA, GH: analyzed data; CMR: conceptualized experimental design, reviewed and edited manuscript drafts and acquired funding.

## Competing Interest Statement

The authors declare no competing interest.

---

**Supplementary Figure 1. Optimization of fluorescence imaging parameters to quantify viability of *Burkholderia glumae* (*pGFP*) cells.** Untreated and heat-killed *B. glumae* (*pGFP*) cells stained with propidium iodide (PI) to differentiate live cells (green fluorescence) from dead cells (red fluorescence) by confocal microscopy (A) and flow cytometry (B).

**Supplementary Figure 2. The *Pseudomonas protegens* PBL3 secretome alters *B. glumae* size, morphology, and envelope ultrastructure. (A)** Flow cytometry average forward-scatter light was used as an indicator of relative cell size of *B. glumae* grown in medium amended with M9 minimal medium or with 0.25 g/mL *P. protegens* PBL3 secretome for 8 and 18 h. (**B**) Scanning electron micrographs of *B. glumae* grown in medium amended with M9 minimal medium or with 0.25 g/mL *P. protegens* PBL3 secretome for 18 h. Secretome-treated cells exhibited abundant cell debris (red arrows) and crevices in the cells (yellow arrows) compared to control. Scale bars, 2 µm. (C)

**Supplementary Figure 3. Global transcriptional changes in *Burkholderia glumae* induced by the *Pseudomonas protegens* PBL3 secretome. (A)** Principal component analysis (PCA) of RNA-seq transcriptomic profiles from *B. glumae* treated with the *P. protegens* PBL3 secretome or M9 minimal medium (control) at 4 or 8 h after treatment. Each point represents an independent biological replicate (n = 3). **(B)** Volcano plots showing differential gene expression in *B. glumae* treated with the PBL3 secretome relative to the corresponding M9-treated controls at 4 h and 8 h after treatment. Each point represents an individual gene. Significantly upregulated and downregulated genes are shown in red and blue, respectively, whereas non significantly altered genes are shown in gray. Vertical and horizontal dashed lines indicate differential-expression thresholds of |log₂ fold change| > 1 and Benjamini–Hochberg-adjusted *p* value < 0.05, respectively. Differential expression was determined using DESeq2.

## Notes

### Competing Interest Statement

The authors have declared no competing interest.

## References

Andrews, S. (2010). FastQC: a quality control tool for high throughput sequence data. http://www.bioinformatics.babraham.ac.uk/projects/fastqc

Bajgelman, M. C. (2019). Principles and applications of flow cytometry. In G. Misra, ed (Ed.), Data Processing Handbook for Complex Biological Data Sources (pp. 119–124).

Bolger, A. M., Lohse, M., & Usadel, B. (2014). Trimmomatic: a flexible trimmer for Illumina sequence data. Bioinformatics, 30(15), 2114–2120. 10.1093/bioinformatics/btu170

Butler, L., Awan, A. R., Ellis, T., & Akram, M. S. (2026). Engineering non-ribosomal peptide synthesis: tuning the antibiotics engine of the microbial world. Crit Rev Biotechnol, 46(3), 501–521. 10.1080/07388551.2026.2615819

Dahal, S., Alvarez, S., Balboa, S. J., Hicks, L. M., & Rojas, C. M. (2024). Defining the Secondary Metabolites in the *Pseudomonas protegens* PBL3 Secretome with Antagonistic Activity Against *Burkholderia glumae*. Phytopathology, 114(12), 2481–2490. 10.1094/PHYTO-04-24-0140-R

Dahal, S., Grosz, K., & Rojas, C. M. (2025). Advancing the Development of Pseudomonas protegens PBL3 Secretome as a Biopesticide to Control Bacterial Panicle Blight in Rice. Plant Pathology, 75(1). 10.1111/ppa.70102

Davey, H., & Guyot, S. (2020). Estimation of Microbial Viability Using Flow Cytometry. Curr Protoc Cytom, 93(1), e72. 10.1002/cpcy.72

Delcour, A. H. (2009). Outer membrane permeability and antibiotic resistance. Biochim Biophys Acta, 1794(5), 808–816. 10.1016/j.bbapap.2008.11.005

Feng, J., Zheng, Y., Ma, W., Ihsan, A., Hao, H., Cheng, G., & Wang, X. (2023). Multitarget antibacterial drugs: An effective strategy to combat bacterial resistance. Pharmacol Ther, 108550. 10.1016/j.pharmthera.2023.108550

Fenn, K. L., Horne, J. E., Crossley, J. A., Bohringer, N., Horne, R. J., Schaberle, T. F., Calabrese, A. N., Radford, S. E., & Ranson, N. A. (2024). Outer membrane protein assembly mediated by BAM-SurA complexes. Nat Commun, 15(1), 7612. 10.1038/s41467-024-51358-x

Gray, D. A., & Wenzel, M. (2020). Multitarget Approaches against Multiresistant Superbugs. ACS Infect Dis, 6(6), 1346–1365. 10.1021/acsinfecdis.0c00001

Gu, Q., Qiao, J., Wang, R., Lu, J., Wang, Z., Li, P., Zhang, L., Ali, Q., Khan, A. R., Gao, X., & Wu, H. (2022). The Role of Pyoluteorin from *Pseudomonas protegens* Pf-5 in Suppressing the Growth and Pathogenicity of *Pantoea ananatis* on Maize. Int J Mol Sci, 23(12). 10.3390/ijms23126431

Halawa, E. M., Fadel, M., Al-Rabia, M. W., Behairy, A., Nouh, N. A., Abdo, M., Olga, R., Fericean, L., Atwa, A. M., El-Nablaway, M., & Abdeen, A. (2023). Antibiotic action and resistance: updated review of mechanisms, spread, influencing factors, and alternative approaches for combating resistance. Front Pharmacol, 14, 1305294. 10.3389/fphar.2023.1305294

Hutchings, M. I., Truman, A. W., & Wilkinson, B. (2019). Antibiotics: past, present and future. Current Opinion in Microbiology, 51, 72–80. 10.1016/j.mib.2019.10.008

Iacovelli, R., Bovenberg, R. A. L., & Driessen, A. J. M. (2021). Nonribosomal peptide synthetases and their biotechnological potential in. Journal of Industrial Microbiology & Biotechnology, 48(7-8). 10.1093/jimb/kuab045

Julian, W. T., Vasilchenko, A. V., Shpindyuk, D. D., Poshvina, D. V., & Vasilchenko, A. S. (2021). Bacterial-Derived Plant Protection Metabolite 2,4-Diacetylphloroglucinol: Effects on Bacterial Cells at Inhibitory and Subinhibitory Concentrations. Biomolecules, 11(1). 10.3390/biom11010013

Katz, L., & Baltz, R. H. (2016). Natural product discovery: past, present, and future. J Ind Microbiol Biotechnol, 43(2-3), 155–176. 10.1007/s10295-015-1723-5

Kim, D., Langmead, B., & Salzberg, S. L. (2015). HISAT: a fast spliced aligner with low memory requirements. Nat Methods, 12(4), 357–360. 10.1038/nmeth.3317

King, E. O., Ward, M. K., & Raney, D. E. (1954). Two simple media for the demonstration of pyocyanin and fluorescin. J Lab Clin Med, 44(2), 301–307. https://www.ncbi.nlm.nih.gov/pubmed/13184240

Kohanski, M. A., Dwyer, D. J., & Collins, J. J. (2010). How antibiotics kill bacteria: from targets to networks. Nature Reviews Microbiology, 8(6), 423–435. 10.1038/nrmicro2333

Li, H., Handsaker, B., Wysoker, A., Fennell, T., Ruan, J., Homer, N., Marth, G., Abecasis, G., Durbin, R., & Genome Project Data Processing, S. (2009). The Sequence Alignment/Map format and SAMtools. Bioinformatics, 25(16), 2078–2079. 10.1093/bioinformatics/btp352

Liao, Y., Smyth, G. K., & Shi, W. (2014). featureCounts: an efficient general purpose program for assigning sequence reads to genomic features. Bioinformatics, 30(7), 923–930. 10.1093/bioinformatics/btt656

Love, M. I., Huber, W., & Anders, S. (2014). Moderated estimation of fold change and dispersion for RNA-seq data with DESeq2. Genome Biol, 15(12), 550. 10.1186/s13059-014-0550-8

Mahone, C. R., & Goley, E. D. (2020). Bacterial cell division at a glance. Journal of Cell Science, 133(7). 10.1242/jcs.237057

Ntallis, C., Martin, N. I., Edwards, A. M., & Weingarth, M. (2026). Bacterial cell envelope-targeting antibiotics. Nature Reviews Microbiology, 24(3), 183–196. 10.1038/s41579-025-01247-x

O’Donnell, M., Langston, L., & Stillman, B. (2013). Principles and concepts of DNA replication in bacteria, archaea, and eukarya. Cold Spring Harb Perspect Biol, 5(7). 10.1101/cshperspect.a010108

Ojha, B., Grosz, K., & Rojas, C. M. (2025). The secretome of the environmental bacterium Pseudomonas protegens PBL3 has broad-spectrum antimicrobial activity against plant and human pathogenic bacteria. *Microbiol Spectr*, e0266925. 10.1128/spectrum.02669-25

Ortega, L., Walker, K. A., Patrick, C., Wamishe, Y., Rojas, A., & Rojas, C. M. (2020). Harnessing *Pseudomonas protegens* to Control Bacterial Panicle Blight of Rice. Phytopathology, 110(10), 1657–1667. 10.1094/PHYTO-02-20-0045-R

Page, J. E., & Walker, S. (2021). Natural products that target the cell envelope. Current Opinion in Microbiology, 61, 16–24. 10.1016/j.mib.2021.02.001

Patil, P. S., & Gore, N. S. (2025). Bacterial non-ribosomal peptides: a sustainable approach for combatting phytopathogenic fungi in agriculture. New Zealand Journal of Botany, 63(5), 1808–1822. 10.1080/0028825x.2024.2401405

Raetz, C. R., Guan, Z., Ingram, B. O., Six, D. A., Song, F., Wang, X., & Zhao, J. (2009). Discovery of new biosynthetic pathways: the lipid A story. J Lipid Res, 50 *Suppl*(Suppl), S103–108. 10.1194/jlr.R800060-JLR200

Reyes-Lamothe, R., & Sherratt, D. J. (2019). The bacterial cell cycle, chromosome inheritance and cell growth. Nature Reviews Microbiology, 17(8), 467–478. 10.1038/s41579-019-0212-7

Reygaert, W. C. (2018). An overview of the antimicrobial resistance mechanisms of bacteria. AIMS Microbiol, 4(3), 482–501. 10.3934/microbiol.2018.3.482

Rojas, C. M., Ham, J. H., Deng, W. L., Doyle, J. J., & Collmer, A. (2002). HecA, a member of a class of adhesins produced by diverse pathogenic bacteria, contributes to the attachment, aggregation, epidermal cell killing, and virulence phenotypes of Erwinia chrysanthemi EC16 on Nicotiana clevelandii seedlings. Proc Natl Acad Sci U S A, 99(20), 13142–13147. 10.1073/pnas.202358699

Schindelin, J., Arganda-Carreras, I., Frise, E., Kaynig, V., Longair, M., Pietzsch, T., Preibisch, S., Rueden, C., Saalfeld, S., Schmid, B., Tinevez, J. Y., White, D. J., Hartenstein, V., Eliceiri, K., Tomancak, P., & Cardona, A. (2012). Fiji: an open-source platform for biological-image analysis. Nat Methods, 9(7), 676–682. 10.1038/nmeth.2019

Silhavy, T. J., Kahne, D., & Walker, S. (2010). The bacterial cell envelope. Cold Spring Harb Perspect Biol, 2(5), a000414. 10.1101/cshperspect.a000414

Silver, L. L. (2007). Multi-targeting by monotherapeutic antibacterials. Nat Rev Drug Discov, 6(1), 41–55. 10.1038/nrd2202

Talapko, J., Mestrovic, T., Juzbasic, M., Tomas, M., Eric, S., Horvat Aleksijevic, L., Bekic, S., Schwarz, D., Matic, S., Neuberg, M., & Skrlec, I. (2022). Antimicrobial Peptides-Mechanisms of Action, Antimicrobial Effects and Clinical Applications. Antibiotics (Basel), 11(10). 10.3390/antibiotics11101417

Wenzel, M. (2020). Do we really understand how antibiotics work? Future Microbiology, 15(14), 1307–1311. 10.2217/fmb-2019-0324

