## Supplementary figures and images for "The *Pseudomonas protegens* PBL3 secretome exerts antimicrobial activity against *Burkholderia glumae* by disrupting cell envelope homeostasis and impairing bacterial replication"

### Supplementary Figures

A

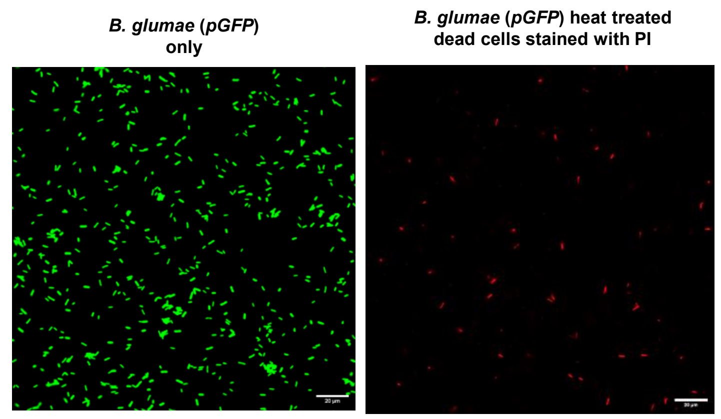


B

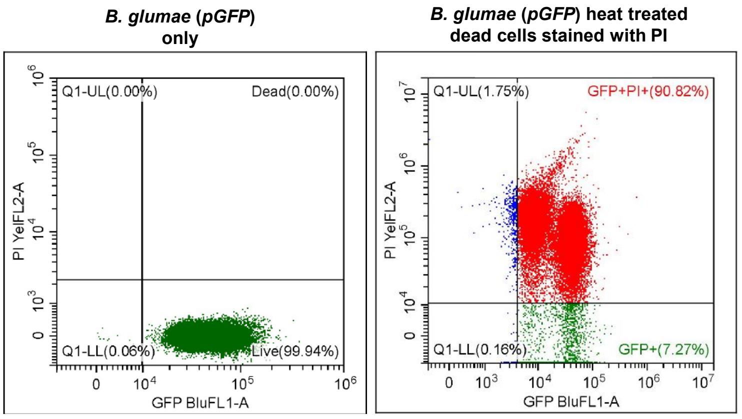


#### Supplementary Figure S1

A

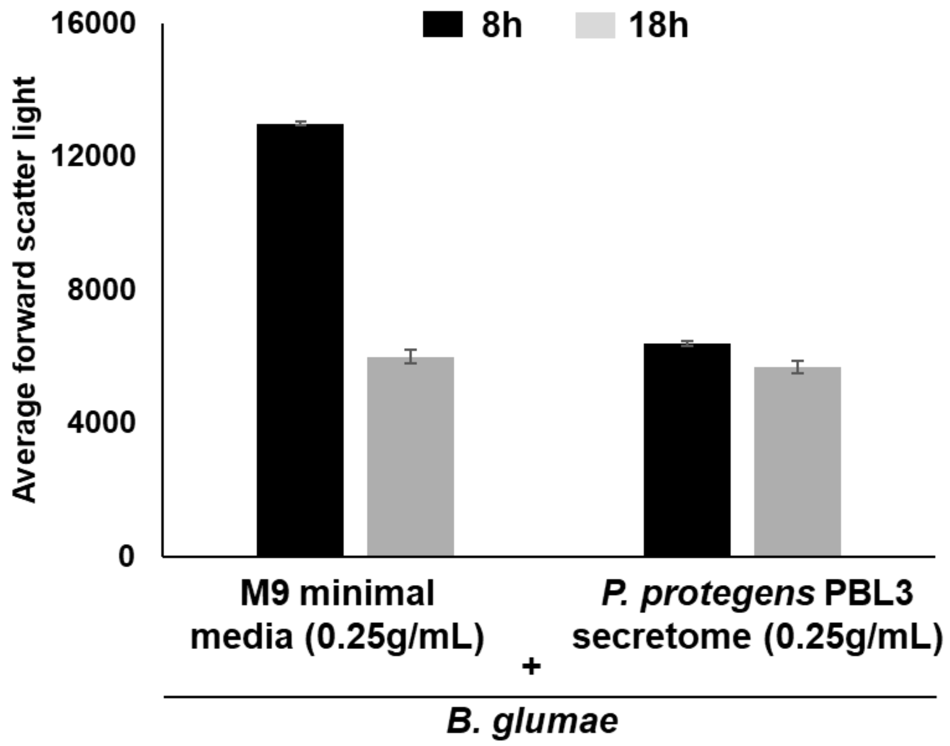


B

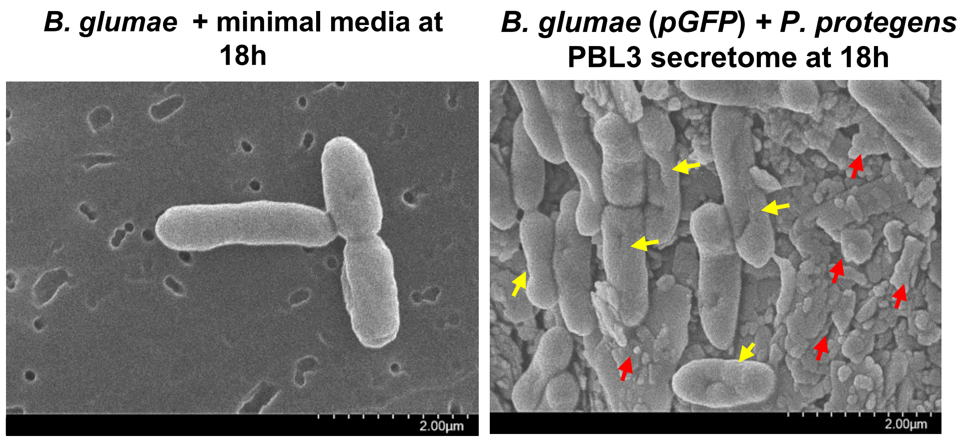


**Supplementary Figure S2**

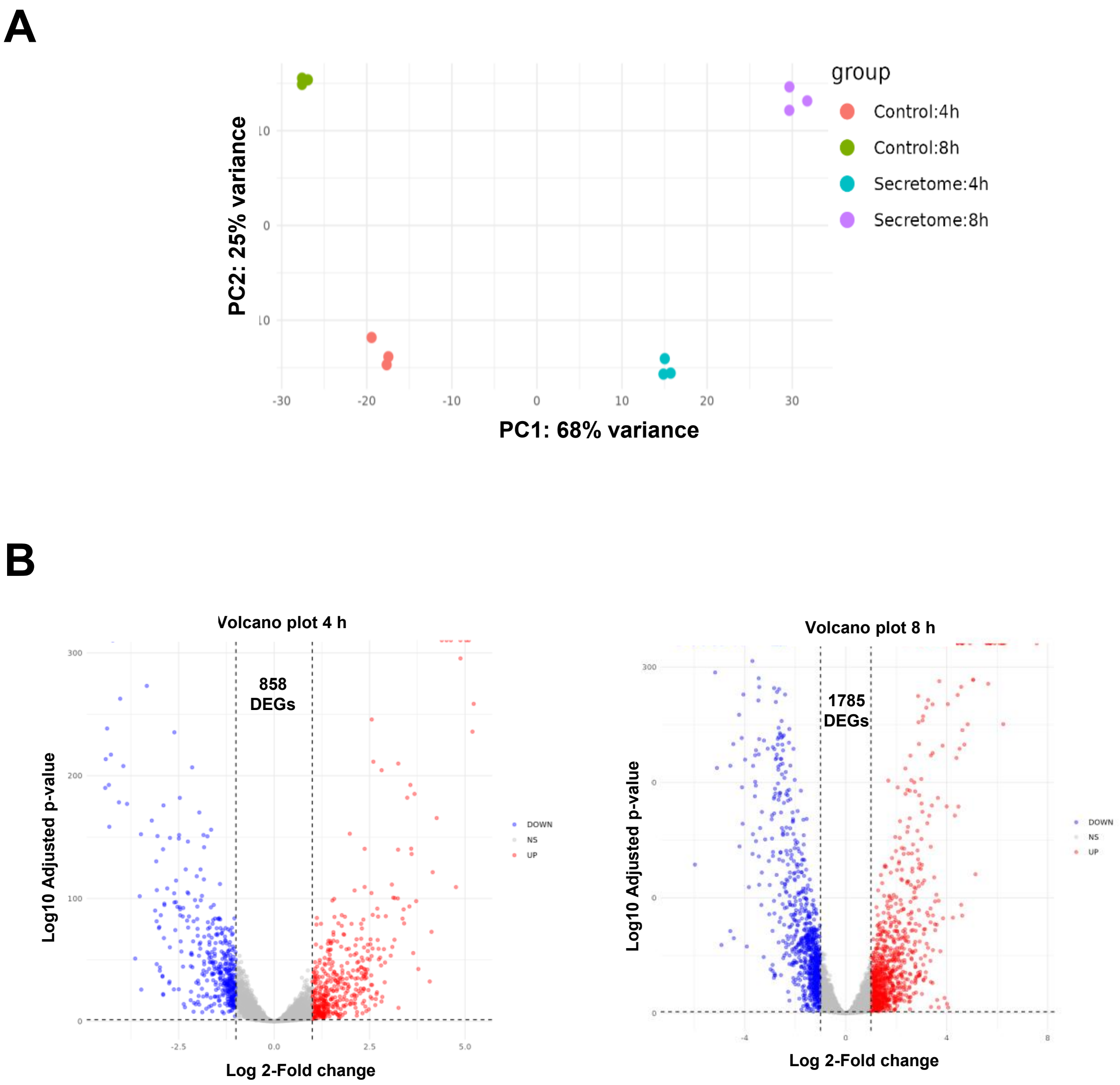


**Supplementary Figure S3**
